# Initiation of chromosome replication controls both division and replication cycles in E. coli through a double-adder mechanism

**DOI:** 10.1101/593590

**Authors:** Guillaume Witz, Erik van Nimwegen, Thomas Julou

## Abstract

Living cells proliferate by completing and coordinating two essential cycles, a division cycle that controls cell size, and a DNA replication cycle that controls the number of chromosomal copies in the cell. Despite lacking dedicated cell cycle control regulators such as cyclins in eukaryotes, bacteria such as *E. coli* manage to tightly coordinate those two cycles across a wide range of growth conditions, including situations where multiple nested rounds of replication progress simultaneously. Various cell cycle models have been proposed to explain this feat, but it has been impossible to validate them so far due to a lack of experimental tools for systematically testing their different predictions. Recently new insights have been gained on the division cycle through the study of the structure of fluctuations in growth, size, and division in individual cells. In particular, it was found that cell size appears to be controlled by an adder mechanism, *i.e.* the added volume between divisions is held approximately constant and fluctuates independently of growth rate and cell size at birth. However, how replication initiation is regulated and coupled to cell size control remains unclear, mainly due to scarcity of experimental measurements on replication initiation at the single-cell level. Here, we used time-lapse microscopy in combination with microfluidics to directly measure growth, division and replication in thousands of single *E. coli* cells growing in both slow and fast growth conditions. In order to compare different phenomenological models of the cell cycle, we introduce a statistical framework which assess their ability to capture the correlation structure observed in the experimental data. Using this in combination with stochastic simulations, our data indicate that, instead of thinking of the cell cycle as running from birth to division, one should consider the chromosome replication cycle as central and in control of the cell cycle through two adder mechanisms: the added volume since the last initiation event controls the timing of both the next division event and the next replication initiation event. Interestingly the double-adder mechanism identified in this study has recently been found to explain the more complex cell cycle of mycobacteria, suggesting shared control strategies across species.

## Introduction

Across all domains of life, cell proliferation requires that the chromosome replication and cell division cycles are coordinated to ensure that every new cell receives one copy of the genetic material. While in eukaryotes this coordination is implemented by a dedicated regulatory system in which genome replication and division occur in well-separated stages, no such system has been found in most bacteria. This suggests that the molecular events that control replication initiation and division might be coordinated more directly in bacteria, through molecular interactions that are yet to be elucidated. The contrast between this efficient coordination and the apparent absence of a dedicated regulatory system is particularly remarkable since most bacteria feature a unique replication origin which imposes that multiple rounds of replication occur concurrently in fast growth conditions. For example, in the specific case of *E. coli* that we study here, it has long been known that growth rate, cell size, and replication initiation are coordinated such that the average number of replication origins per unit of cellular volume is approximately constant across conditions (***Donachie, 1968***) or that cellular volume grows approximately exponentially with growth rate (***Taheri-Araghi et al., 2017***). Although several models have been proposed over the last decades to explain such observations, so far direct validation of these models has been lacking, due to a large extent to the lack of quantitative measurements of cell cycles parameters in large samples with single-cell resolution.

Thanks to techniques such as microfluidics and time-lapse microscopy, it has recently become possible to perform long-term observation of growth and division in single bacteria. By systematically quantifying how cell cycle variables such as size at birth, size at division, division time, and growth rate vary across single cells, insights can be gained about the mechanism of cell cycle control. Several recent studies have focused on understanding the regulation of cell size, resulting in the discovery that *E. coli* cells maintain a constant average size by following an adder strategy: instead of attempting to reach a certain size at division (*i.e.* a sizer mechanism) or to grow for a given time (*i.e.* a timer mechanism), it was found that cells add a constant length *dL* to their birth length *L*_*b*_ before dividing (***Amir, 2014***; ***Campos et al., 2014***; ***Taheri-Araghi et al., 2017***). In particular, while the cell size at division and the division time correlate with other variables such the cell size at birth and growth rate, the added length *dL* fluctuates independently of birth size and growth rate. A remarkable feature of the adder model is its capacity to efficiently dampen large cell size fluctuations caused by the intrinsically noisy regulation, without the need for any fail-safe mechanism. This efficient strategy has been shown to be shared by various bacterial species as well as by archea (***Eun et al., 2018***) and even some eukaryotes such as budding yeast (***Soifer et al., 2016***).

Here we focus on how the control of replication initiation is coordinated with cell size control in *E. coli*. Several models have been proposed to explain how the adder behavior at the level of cell size might arise from a coordinated control of replication and division. Broadly, most models assume that the accumulation of a molecular trigger, usually assumed to be DnaA, leads to replication initiation, which in turn controls the corresponding future division event (***Campos et al., 2014***; ***Ho and Amir, 2015***; ***Wallden et al., 2016***). Subtle variations in how the initiation trigger accumulates and how the initiation to division period is set in each model imply distinct molecular mechanisms, and thus fundamentally different cell cycle regulations. Specifically, most models assume that initiation is triggered either when a cell reaches a critical absolute volume (initiation size, see *e.g.* ***Wallden et al., 2016***) or alternatively when it has accumulated a critical volume since the last initiation event (see *e.g.* ***Ho and Amir, 2015***). In order to explain the coordination between cell cycle events, division is often assumed to be set by a timer starting at replication initiation, but recent studies have also proposed that the two cycles might be independently regulated (***Micali et al., 2018a***; ***Si et al., 2019***). Finally, it is often assumed that the regulation strategy could be different at slow and fast growth where different constraints occur.

We use an integrated microfluidics and time-lapse microscopy approach to quantitatively characterize growth, division, and replication in parallel across many lineages of single *E. coli* cells, both in slow and fast growth conditions. We show that insights about the underlying control mechanisms can be gained by systematically studying the structure of correlations between these different variables. Our single-cell observations are inconsistent with several previously proposed models including models that assume replication is initiated at a critical absolute cell volume and models that assume division is set by a timer that starts at replication initiation. Instead, the most parsimonious model consistent with our data is a double-adder model in which the cell cycle commences at initiation of replication and both the subsequent division and the next initiation of replication are controlled by the added volume. We show that this model is most consistent with the correlation structure of the fluctuations in the data and, through simulations, we show that this model accurately reproduces several non-trivial observables including the previously observed adder behavior for cell size control, the distribution of cell sizes at birth, and the distribution of the number of origins per cell. Moreover, the same model best describes the data both at slow and fast growth rates. As far as we aware, no other proposed model can account for the full set of observations we present here.

## Results

To test possible models for the coordination of replication and division in *E. coli* we decided to systematically quantify growth, replication initiation, and division across thousands of single *E. coli* cell cycles, across multiple generations, and in various growth conditions. To achieve this, cells were grown in a Mother Machine type microfluidic device (***Wang et al., 2010***) and imaged by time-lapse microscopy. We used M9 minimal media supplemented with glycerol, glucose or glucose and 8 amino acids, resulting in doubling times of 89, 53 and 41 min, respectively. The cell growth and division cycles were monitored by measuring single-cell growth curves obtained through segmentation and tracking of cells in phase contrast images using the MoMA software (***Kaiser et al., 2018***). The replication cycle was monitored by detecting initiation as the duplication of an *oriC* proximal FROS tagged *locus* imaged by fluorescence microscopy (***Figure 1A***). These measurements allowed us to quantify each single cell cycle by a number of variables such as the growth rate, the sizes at birth, replication initiation, and division, the times between birth and replication initiation and the time between birth and division. As done previously, we assume that cell radius is constant and use cell length as a proxy for cell volume (***Adiciptaningrum et al., 2015***; ***Taheri-Araghi et al., 2017***). Since we can follow cells over multiple generations, we can also measure quantities that span multiple division cycles such as the total time or total cell growth between consecutive replication initiation events. As we analyze growth conditions spanning cases with both single and overlapping rounds of replication, we defined a consistent way of measuring variables. Noticeably, while the cell cycle is classically defined from division to division (***Figure 1C***), as has been proposed previously ***Ho and Amir (2015***); ***Amir (2017***), we use an alternative framework where the cell cycle is defined from one replication initiation to the next (***Figure 1D***). This framework being centered on origins of replication rather than on cells, we consequently define a new quantity Λ, the cell length per origin, which allows to track the growth allocated to a given origin of replication. For instance, in a case where a cell is born with an ongoing round of replication which started at time *t*, Λ_*i*_ for that cell is defined as Λ_*i*_ = *L*_*i*_/4 where *L*_*i*_ is the length of the mother cell which contains four origins at time *t* (***Figure 1 – Figure Supplement 1***). This avoids artificial cut-offs as *e.g.* done in ***Wallden et al. (2016***). In this article, we explore a series of models belonging to these two views of the cell cycle. Using the correlation structure of variables, we show how classes of models can be rejected entirely. Additionally, we use a more general statistical framework to rank models according to their explanatory power.

**Figure 1.**
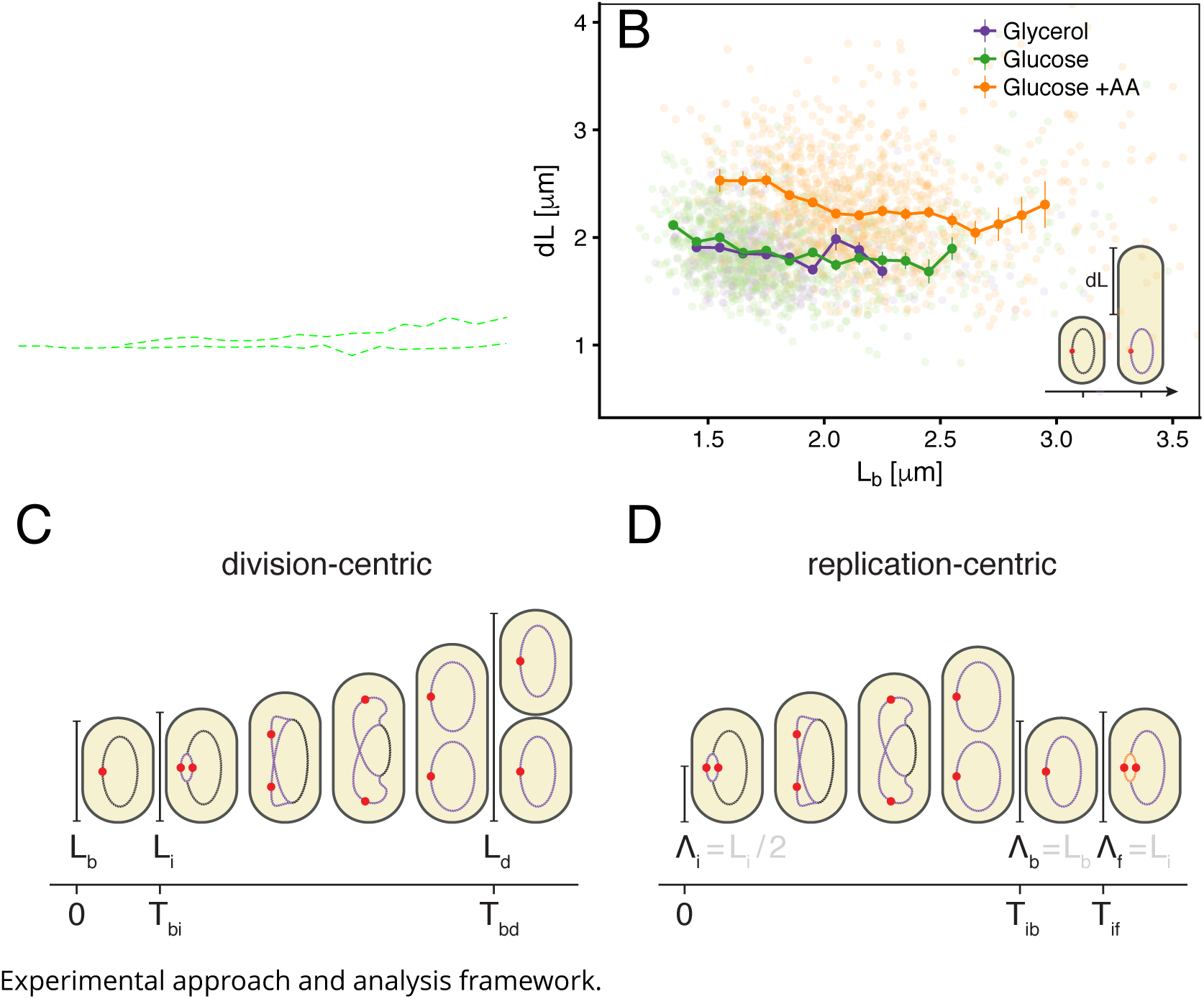
A. Time-lapse of *E. coli* cells growing in a single microfluidic channel. Fluorescence signal from FROS labeling is visible as red spots in each cell. The green dotted line is an aid to the eye, illustrating the replication of a single origin. **B**. Consistent with an adder behavior, the added length between birth and division is uncorrelated with length at birth. **C**. The classical cell cycle is defined between consecutive division events, shown here with replication and division for slow growth conditions (i.e. without overlapping rounds of replication). **D**. We introduce an alternative description framework where the cell cycle is defined between consecutive replication initiation events. The observables that are relevant to characterize the cell cycle in these two frameworks are indicated (see also ***Table 1***). **Figure 1– Figure supplement 1.** Schema of the cell cycle and variable definitions for the case of fast growth with overlapping replication cycles.

### Cell size adder

We first verified whether our measurements support the previously observed adder behavior in cell size, and find that added length *dL* between birth and division is indeed uncorrelated with length at birth *L*_*b*_ in all growth conditions (***Figure 1B***), and, also in agreement with the adder model, the heritability of birth length between mother and daughter is characterized by a Pearson correlation coefficient of *r* ≈ 0.5 (see Table 1). With the exception of one study (***Wallden et al., 2016***), moderately slow growth conditions (100 min doubling time) have not been yet tested extensively for adder behavior. The fact that we observe it in conditions where replication occurs both in overlapping- and non-overlapping modes further highlights its pervasiveness.

**Table 1.** Variables definitions.

| division-centric |  | replication-centric |  |
| --- | --- | --- | --- |
| *measured variables* |  |  |  |
| $L_b$ | Size at birth* | $\Lambda_i$ | Size per origin at *initial* replication initiation* |
| $L_d$ | Size at division* | $\Lambda_f$ | Size per origin at *final* replication initiation* |
| $T_{bd}$ | Duration between birth and division | $T_{if}$ | Duration between consecutive replication initiations |
| $L_i$ | Size at replication initiation* | $\Lambda_b$ | Size per origin at birth* |
| $T_{bi}$ | Duration between birth and replication initiation | $T_{ib}$ | Duration between replication initiation and birth |
| *derived variables* |  |  |  |
| $\lambda = \frac{1}{T_{bd}} \log \frac{L_d}{L_b}$ | Cell growth rate* (between birth and division) | $\alpha = \frac{1}{T_{if}} \log \frac{\Lambda_f}{\Lambda_i}$ | Cell growth rate* (between consecutive replication initiations) |
| $dL = L_d - L_b$ | Division “adder” | $d\Lambda_{if} = \Lambda_f - \Lambda_i$ | Replication “adder” |
| $dL_{bi} = L_i - L_b$ | Birth-to-initiation “adder” | $d\Lambda_{ib} = \Lambda_b - \Lambda_i$ | Initiation-to-birth “adder” |
| $R_{bd} = L_d/L_b$ | Growth ratio between birth and division | $R_{if} = \Lambda_f/\Lambda_i$ | Growth ratio between consecutive initiations |
| $R_{bi} = L_i/L_b$ | Growth ratio between birth and initiation | $R_{ib} = \Lambda_b/\Lambda_i$ | Growth ratio between initiation and birth |

\* variables indicated by a star are measured from a linear fit of exponential elongation.

### Replication initiation mass

A popular idea dating back to the 1960’s and still often used today to explain the coupling of division and replication cycles is the initiation mass model. The observations that cell volume grows exponentially with growth rate (***Schaechter et al., 1958***) and that, across a range of conditions, the time between replication initiation and division is roughly constant (***Helmstetter et al., 1968***) led Donachie to propose that the volume per origin of replication is held constant (***Donachie, 1968***). In particular, the model proposes that initiation occurs when a cell reaches a critical volume. A simple prediction of this model is that, for a given cell, the cell length *L*_*i*_ at which initiation occurs should be independent of other cell cycle variables such as the length at birth *L*_*b*_. However, as can be seen in ***Figure 2A***, we observe that the initiation length *L*_*i*_ and birth length *L*_*b*_ are clearly correlated in all conditions, rejecting the initiation mass model. The absence of an initiation mass has been noted recently elsewhere (***Micali et al., 2018a***). It should be noted, however, that even though the single-cell fluctuations show that initiation is unlikely to be triggered by a critical volume per origin, the constancy of the average volume per origin at initiation across growth conditions (***Si et al., 2017***) still indicates that initiation is probably regulated through a measure of cell volume.

**Figure 2.**
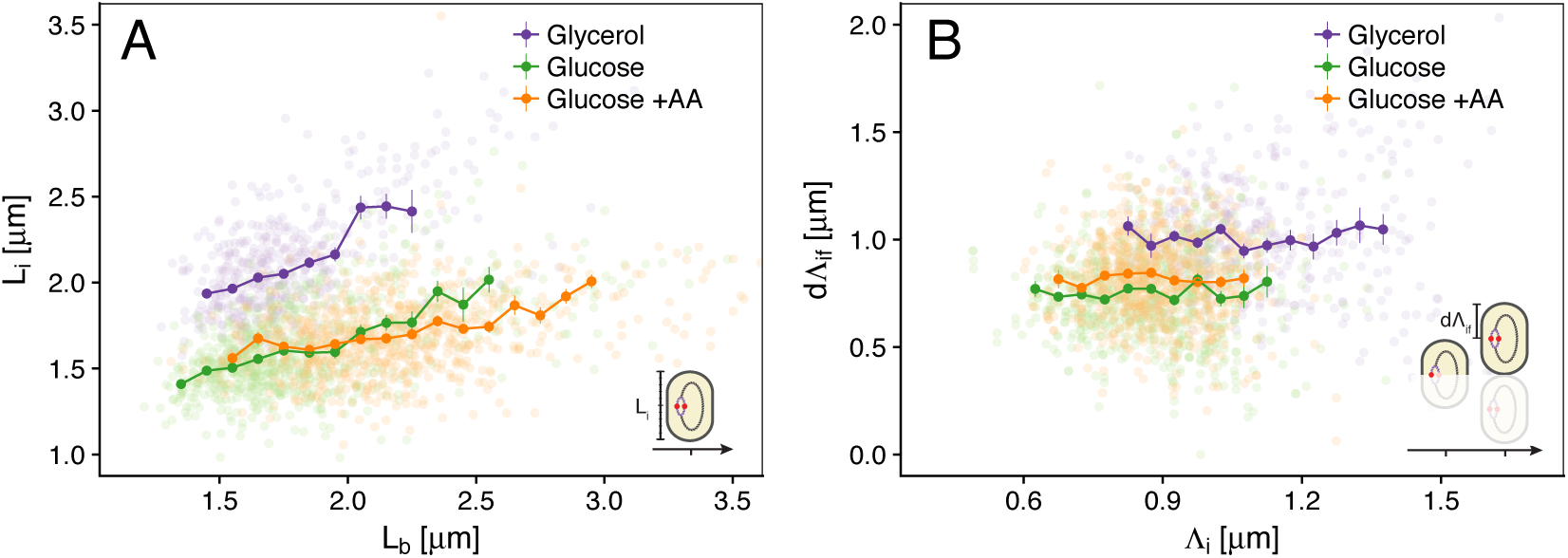
Models for initiation control. **A**. The initiation mass model predicts that the length at initiation *L*_*i*_ should be independent of the length at birth *L*_*b*_. However, we observe clear positive correlations between *L*_*i*_ and *L*_*b*_ in all growth conditions. **B**. In contrast, the length accumulated between two rounds of replication *d*Λ_*if*_ is independent of the initiation size Λ_*i*_, suggesting that replication initiation may be controlled by an adder mechanism.

### Multiple origins accumulation model

Just as a constant average cell size can be accomplished by adding a constant volume per division cycle rather than by dividing at a critical division volume, so a constant average volume per origin of replication can also be implemented by controlling the added volume between replication initiations rather than by a critical initiation volume. A concrete proposal for such an adder mechanism, called the multiple origins accumulation model, has recently received increasing attention (***Ho and Amir, 2015***). In this model, a molecule that is expressed at a constant cellular concentration accumulates at each origin until it reaches a critical amount, triggering replication, after which it is degraded and starts a new accumulation cycle. Given that, for a molecule at constant concentration, the added volume over some time period is proportional to the amount produced of the molecule, the result of this process is that the cell adds a constant volume per origin *d*Λ_*if*_ between initiation events (with *d*Λ_*if*_ = Λ_*f*_ − Λ_*i*_ where indexes stand for “initial” and “final” respectively, see ***Figure 1D*** and ***Table 1*** for more details). If replication is indeed triggered by such an adder mechanism, then one would expect the observed added lengths *d*Λ_*if*_ to be independent of the length Λ_*i*_ at the previous initiation. As shown in ***Figure 2B***, our data support this prediction.

### Connecting replication and division cycles

Having validated the multiple origins accumulation model for replication control, we now investigate its relation to the division cycle. A common assumption is that the period *T*_*id*_ from initiation to division (classically split into the replication period C and the end of replication to division period D) is constant and independent of growth rate (***Cooper and Helmstetter, 1968***; ***Ho and Amir, 2015***). As visible in ***Figure 3A***, while on average *T*_*id*_ is indeed rather constant across growth conditions, within each condition fast growing cells clearly complete this period faster than slow growing cells. One way to model this behavior is to define an empirical relation between growth rate *λ* and *T*_*id*_ (***Wallden et al., 2016***). However, ***Figure 3B*** reveals another and arguably simpler solution. We find that *d*Λ_*id*_ = Λ_*d*_ − Λ_*i*_, the length per origin added by a cell between initiation and division, has an adder behavior as well: independently of its size at initiation *L*_*i*_, a cell will complete the corresponding division cycle after having accumulated a constant volume per origin *d*Λ_*id*_.

**Figure 3.**
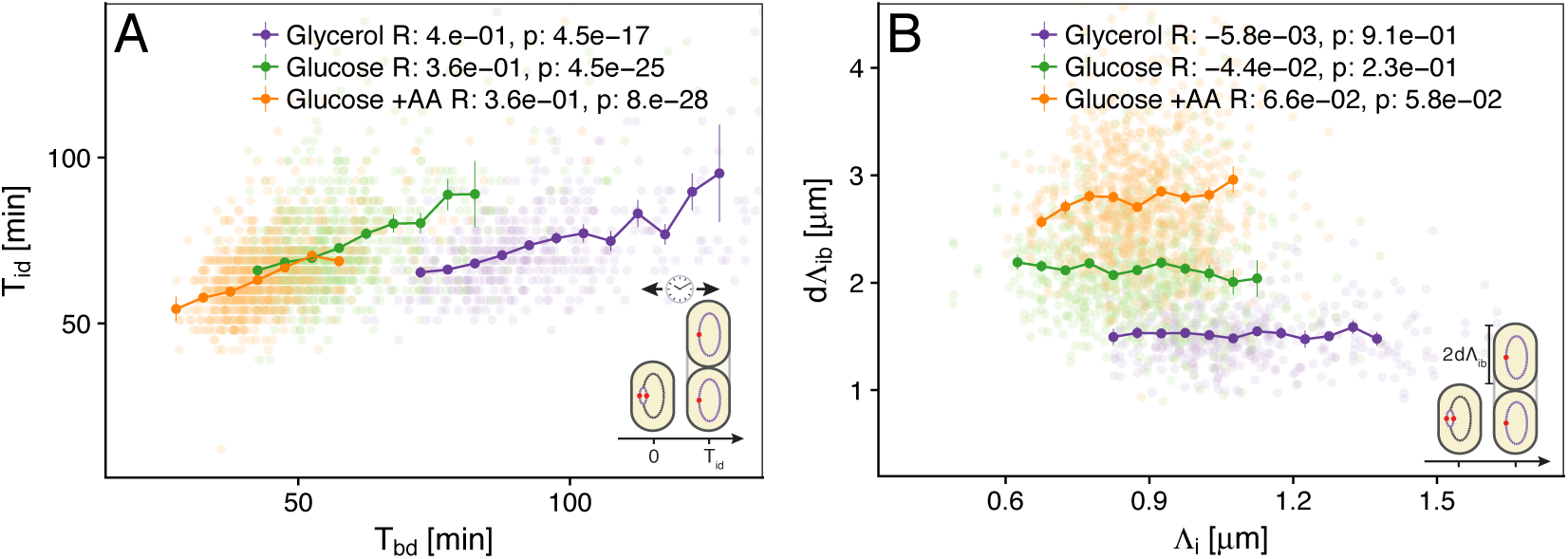
Initiation to division period. A. Several models assume that a constant time passes from an initiation event to it corresponding division event. However, within each growth condition, that period is clearly dependent on fluctuations in growth rate. B. The length accumulated from initiation to division is constant for each growth condition, suggesting an adder behavior for that period. In A and B, the Pearson correlation coefficient R and p values are indicated for each condition.

### The double-adder model

These observations motivated us to formulate a model in which the cell cycle does not run from one division to the next, but rather starts at initiation of replication, and that both the next initiation of replication and the intervening division event, are controlled by two distinct adder mechanisms. In this replication-centric view, the cell cycles are controlled in a given condition by three variables: an average growth rate *λ*, an average added length per origin *d*Λ_*if*_, and an average added length *d*Λ_*id*_ between replication initiation and division. In particular, we assume that these three variables fluctuate independently around these averages for each individual cell cycle, and that all other parameters such as the sizes at birth, initiation, and the times between birth and division or between initiation and division, are all a function of these three fundamental variables. This double-adder model is sketched in ***Figure 4*** for the case of slow growth conditions: a cell growing at a rate *λ* and of length *L* initiates replication and thereby starts two adder processes. First, the cell will divide when reaching a size *n*Λ_*d*_ = *L* + *nd*Λ_*id*_ = *n*(Λ_*i*_ + *d*Λ_*id*_) where *n* = 2 is the number of replication origins. Second, the next replication round will be initiated at a given origin after the corresponding Λ has increased by *d*Λ_*if*_.

**Figure 4.**
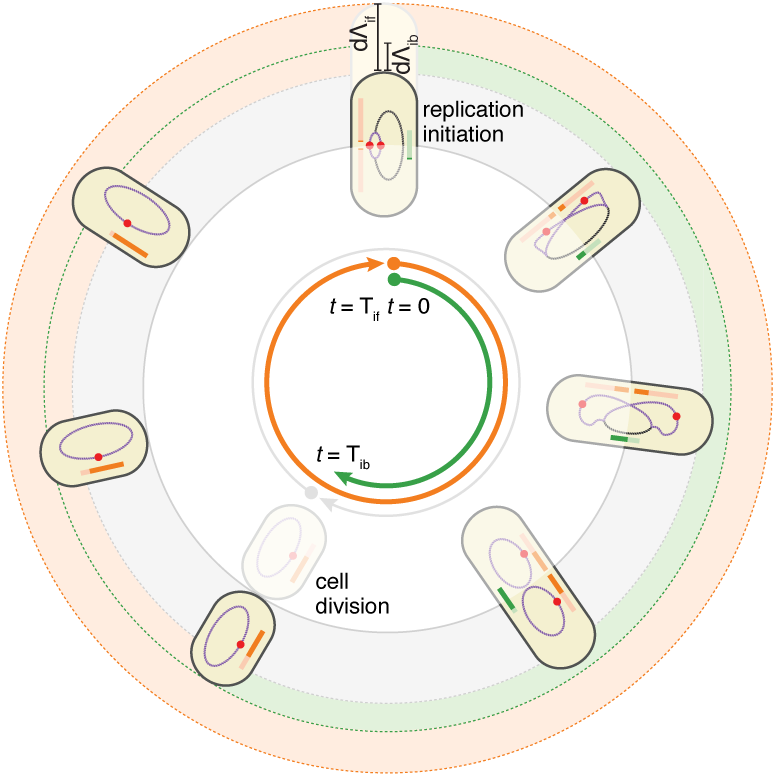
The double-adder model postulates that *E. coli* cell cycle is orchestrated by two independent adders, one for replication and one for division, reset at replication initiation. Both adders (shown as coloured bars) start one copy per origin at replication initiation and accumulate in parallel for some time. After the division adder (green) has reached its threshold, the cell divides, and the initiation adder (orange) splits between the daughters. It keeps accumulating until it reaches its own threshold and initiates a new round of division and replication adders. Note that the double-adder model is illustrated here for the simpler case of slow growth. **Figure 4– Figure supplement 1.** Average localization of the origin in cells growing in M9 glycerol.

### Simulations of the double-adder model

To assess to what extent our double-adder model can recover our quantitative observations, we resorted to numerical simulations. We first obtained from experimental data the empirical distributions of growth rates *λ*, the added length per initiation *d*Λ_*if*_, and the added length between initiation and division *d*Λ_*id*_. A series of cells are initialized at the initiation of replication, with sizes taken from the experimental distributions. For each cell, a growth rate *λ* is independently drawn from its empirical distribution, and values of *d*Λ_*id*_ and *d*Λ_*if*_ are drawn from independent distributions, to set the times of the next division and replication initiation events. This procedure is then iterated indefinitely, *i.e* a new growth rate and values of each adder are independently drawn for each subsequent cycle. As has been observed previously (***Campos et al., 2014***) the growth rate is correlated (*r* ≈ 0.3) between mother and daughter. Accounting for this mother-daughter correlation in growth rate was found not to be critical for capturing features of *E. coli* cell cycle, but was included in the model to reproduce simulation conditions of of previous studies.

As can be seen in ***Figure 5***, the double-adder model accurately reproduces measured distributions and correlations at all growth rates. In particular, the global adder behavior for cell size regulation naturally emerges from it (***Figure 5A***). Similarly, the specific relation between length at initiation *L*_*i*_ and length at birth *L*_*b*_, which prompted us to reject the initiation mass model, is reproduced by the model as well (***Figure 5B***). Finally, the distribution of the number of origins at birth, which reflects the presence of overlapping replication cycles is reproduced as well (***Figure 5D***). An exhaustive comparison between experiments and simulations can be found in ***Figure 5 – Figure Supplement 5***.

**Figure 5.**
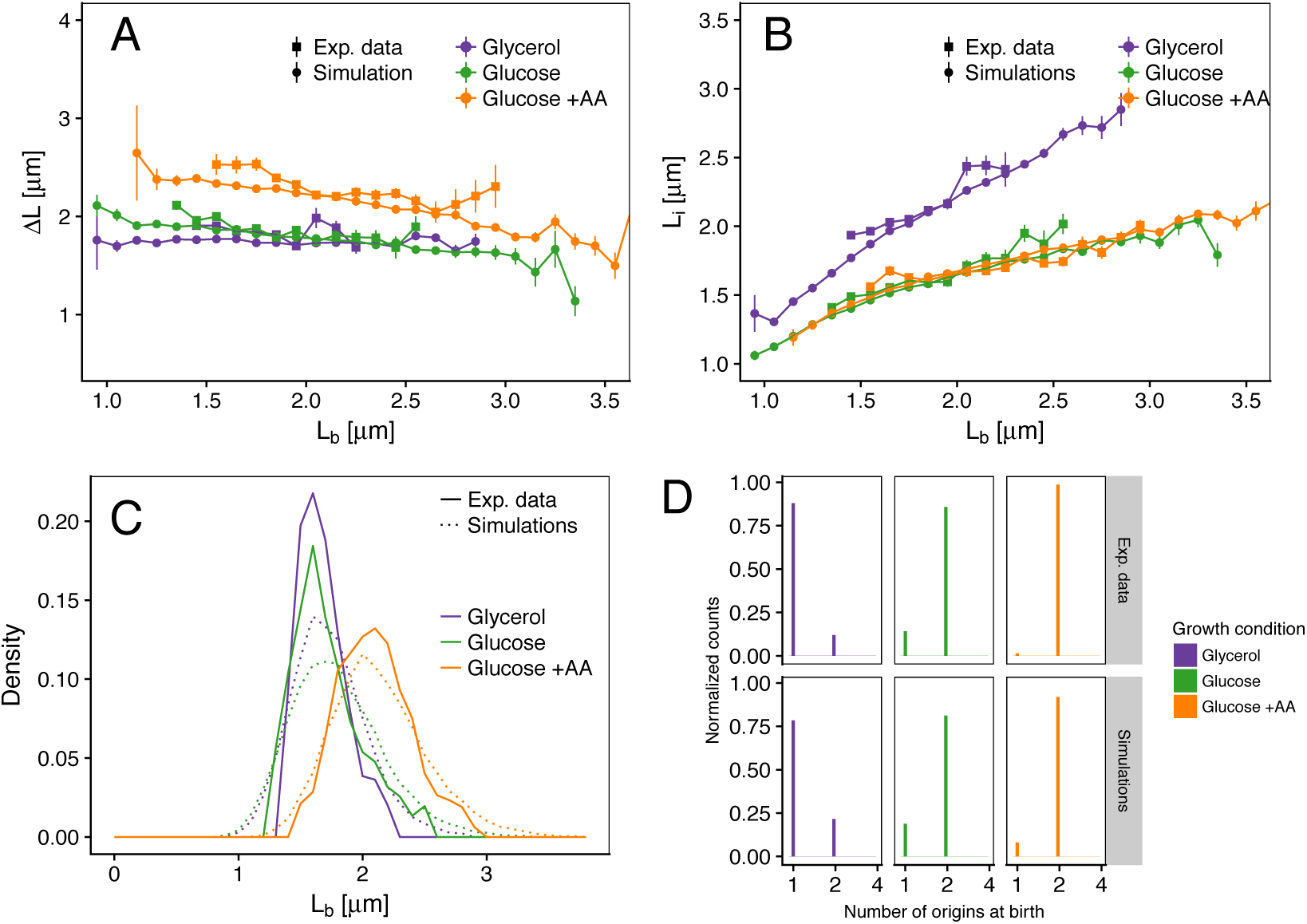
Comparison of predictions of the double-adder model with experimental observations. (A) Added length between birth and division *dL* versus length at birth *L*_*b*_ show no correlations in both the data and the simulations, demonstrating that the double-adder model reproduces the adder behavior at the level of cell size. (B) Length at initiation versus length at birth show almost identical correlations in data and simulation. (C) The distribution of cell sizes at birth are highly similar in experiments (solid lines) and simulations (dashed lines), in all growth conditions. (D) The distribution of the number of origins at birth is also highly similar between experiments and data for all growth conditions. **Figure 5– Figure supplement 1.** Detailed comparisons between experiments and simulations for M9+glycerol condition (with automated origin tracking). **Figure 5– Figure supplement 2.** Detailed comparisons between experiments and simulations for M9+glycerol condition (with manual origin tracking). **Figure 5– Figure supplement 3.** Detailed comparisons between experiments and simulations for M9+glucose condition (with manual origin tracking). **Figure 5– Figure supplement 4.** Detailed comparisons between experiments and simulations for M9+glucose+8a.a. condition (with manual origin tracking). **Figure 5– Figure supplement 5.** Improved simulation by reducing variance.

### The double-adder model best captures the correlation structure of the data

Although our simulations show that the double-adder model, which takes *λ, d*Λ_*id*_ and *d*Λ_*if*_ as the key independently fluctuating quantities, can accurately reproduce our observations, it is less clear whether there are not many other models that could reproduce the data equally well? As the space of possible models is arguably unlimited, it is difficult to answer this question in full generality. However, we can rigorously compare a large class of possible models, by quantitatively comparing the correlation structure that each model implies, with the correlation structure evident in the data. For example, as noted above, the main argument in favor of a cell size adder model is that, whereas birth and division size generally correlate, added volume does not correlate with birth size. Similarly, while the time between birth and division correlates with both the added volume and the growth rate, growth rate and added volume do not correlate. That is, the evidence in favor of a given model can be quantified by the extent to which the key variables of the model are fluctuating independently.

The quantities that are measured directly for each cell cycle are the times and cell sizes at which various events take place. If we take a division-centric view, *i.e.* thinking of each cell cycle as running from birth to division, each cell cycle is characterized by four directly measured quantities: the sizes at birth *L*_*b*_, initiation *L*_*i*_, and division *L*_*d*_, and the doubling time *T*_*bd*_. Similarly, for a replication-centric view, the four directly measured quantities are the sizes per origin at initiation Λ_*i*_, at birth after the subsequent division Λ_*b*_, and at the next initiation Λ_*f*_, as well as the time *T*_*if*_ between consecutive initiations (Fig.1 C-F). However, these directly measured quantities are highly correlated. The correlation structure of the data is captured by the covariance matrix *C*, with diagonal components *C*_*xx*_ corresponding to the variances *V*_*x*_ of each variable *x*, and the off-diagonal components *C*_*xy*_ corresponding to the covariances between pairs of variables (*x, y*). If one thinks of the collection statistics of all single cell cycles as a scatter in 4-dimensional space, then the determinant of the covariance matrix *D*(*C*) can be thought of as the square of the *volume* covered by this 4-dimensional scatter. This squared-volume *D*(*C*) can be at most as large as the product of the variances of all variables *D*(*C*) ≤ *V*_max_ = ∏ _*x*_ *V*_*x*_ with equality if and only if all variables are independent (see Fig.6B for an illustration of the 2D case). That is, the smaller the ratio *D*(*C*)/*V*_max_, the stronger are the correlations of the variables. We call this ratio independence and denote it as *I* = *D*(*C*)/*V*_max_. In ***Appendix 3***, we apply this approach to the simpler case of the sole division cycle that is defined by only three variables. We show that the variables of the adder model constitute the set for which fluctuations are most independent; remarkably *I* ≈ 1 indicates almost full independence in this case.

We can now systematically explore which set of variables best explains the correlation structure in the data by searching for the set of variables that maximizes independence *I*. For example, while a model that assumes a timer between initiation and division would treat the time *T*_*ib*_ as an independent variable, in our double-adder model the variables are the growth rate *λ* and added length *d*Λ_*id*_; by definition, the time *T*_*ib*_ is related to these through 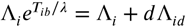. In this way, we can systematically explore different models by taking different sets of variables as fundamental and calculate the independence of each parameter set. Such a statistical analysis is only relevant when applied to a large dataset and we therefore focus here on the slow growth condition (M9 glycerol) for which we implemented automatic origin tracking.

The tables ***Figure 6A*** bottom show the five best models ranked by decreasing independence (all decompositions can be found in ***Figure 6 – Figure Supplement 6***). Note that these variable sets include all the previously proposed sizer and timer models as special cases, for example the inter-initiation model combined with an initiation to division timer is highlighted in red in ***Figure 6 – Figure Supplement 6***. The most successful models are shown in greater detail as correlation matrices Fig.6A top, where residual correlations between all pairs of variables are visible. We find that none of the division-centric models accomplishes high independence. For example, as shown in the correlation matrix, the best division-centric model is plagued by high correlation between *L*_*b*_ and *dL*_*bi*_. This strongly suggests that the cell cycle control is better described from a replication-centric point of view. Of all replication-centric models, our double-adder model clearly reaches the highest independence, followed by various derivative models in which one of the adders is replaced by another variable. We note that independence of our double-adder model on the real data is only slightly lower than on simulated data ***Figure 6D***, *i.e* 0.86 versus 0.98. This residual dependence might either result from correlated errors in the measurements, or it might reflect some small biological dependence not captured by our model. In summary, a systematic analysis shows that, within a large class of alternative models, the double-adder model best captures the correlation structure evident in the data.

**Figure 6.**
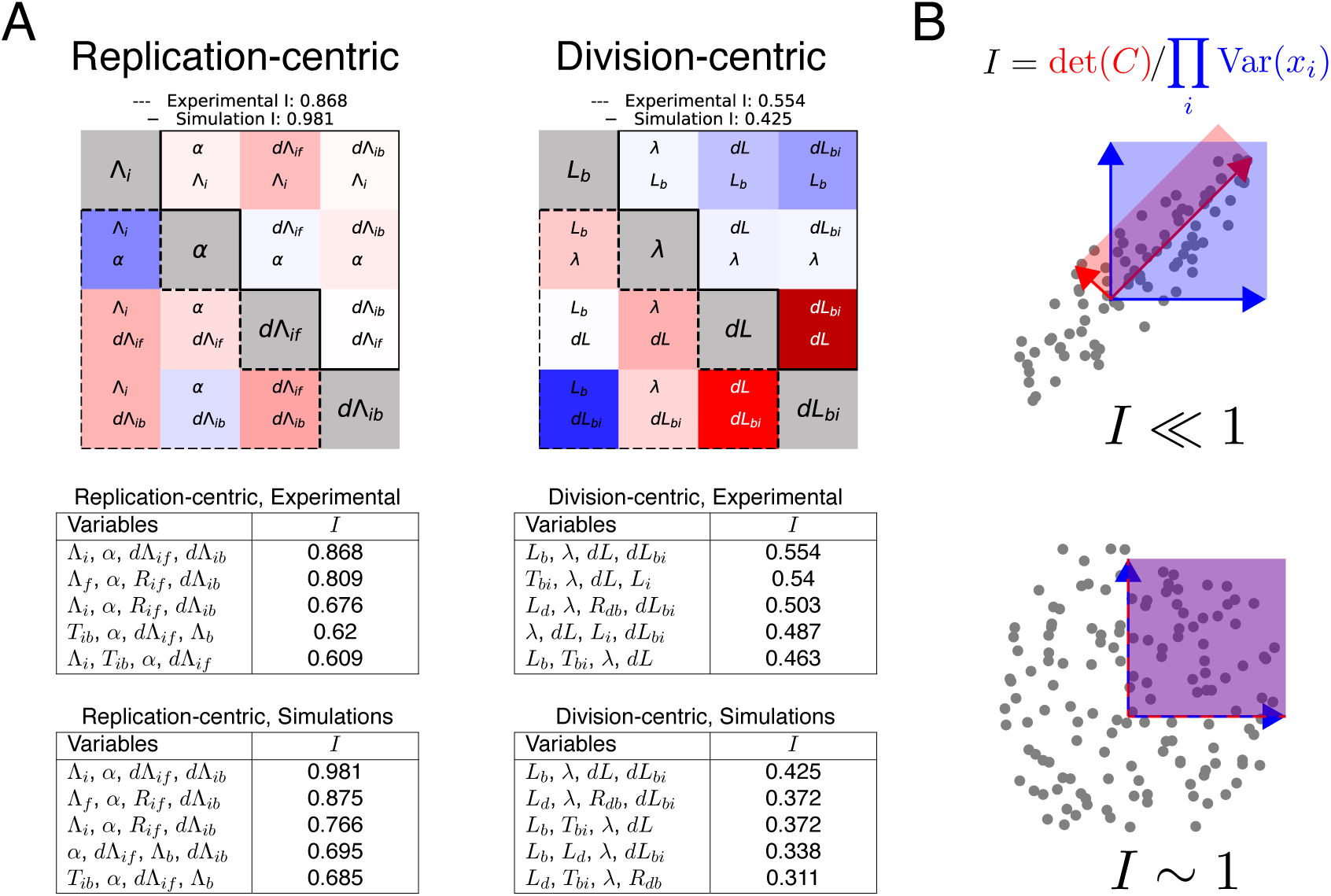
Decomposition method. A. Best decompositions for replication-centric and division-centric models. The matrices show the correlation structure between the variables composing the most independent set of variables. Each square corresponds to a pair of variables, red indicating positive and blue negative correlation. The lower-left corners contain experimental data and the upper-right ones simulation data B. 2D illustration of the independence quantification. The distribution of a pair of variables is shown. The variance of each variable is indicated in blue. The shaded blue area corresponds to the product of variances. The determinant of the correlation matrix of two variables gives the area spanned by the eigenvectors of the matrix (red). The variances are the same for the example with correlations (top) and without (bottom). The area determined by *det*(*C*) is strongly reduced in the correlated case. D. Five best decompositions for replication- and division-centric models of experimental and simulation data. **Figure 6– Figure supplement 1.** Detailed results of the nine best decompositions for experimental data. **Figure 6– Figure supplement 2.** Detailed results of the nine best decompositions for simulation data. **Figure 6– Figure supplement 3.** Complete tables of decomposition.

## Discussion

Thanks to experimental techniques like the one used here, models of bacterial cell cycle regulation dating back from the 1960’s have been recently re-examined in detail in several studies ***Campos et al. (2014***); ***Tanouchi et al. (2015***); ***Ho and Amir (2015***); ***Adiciptaningrum et al. (2015***); ***Wallden et al. (2016***); ***Si et al. (2017***); ***Logsdon et al. (2017***); ***Micali et al. (2018a***); ***Eun et al. (2018***); ***Si et al. (2019***). As much as these new data have been useful in shedding light on regulation mechanisms of bacterial physiology such as the adder, they have also revealed that multiple models are capable of reproducing in large parts experimental data, mainly as a consequence of the correlations existing between measurable cell cycle parameters. However, all of them fail at reproducing at least one important aspect of the experimental observations. To illustrate this, we show in ***Appendix 2*** how two recently proposed models (***Wallden et al., 2016***; ***Micali et al., 2018b***), despite being very successful, fail to capture at least one important observation. In this study, we have in a first step empirically built a model which is based on previous ideas and which recapitulates measured cell cycle parameters. This model makes replication the central regulator of the cell cycle, with each initiation round triggering subsequent division and replication events through an adder mechanism. In a second step, we then constructed and applied a statistical method to determine, within a class of models, which set of cell cycle parameters best explains measurements. Following this fully independent approach, our empirical double-adder model clearly comes out as the most successful.

While the division and replication cycles are seemingly coupled, our analysis demonstrates that two simple adders connected to replication initiation are sufficient to recapitulate both cycles without explicitly enforcing constraints reflecting mechanisms such as over-initiation control by SeqA and nucleoid occlusion which ensures that division only occurs after chromosome replication is completed. The initiation adder mimicks SeqA activity by creating a refractory period without initiation, and the division adder ensures that a minimal time is allocated for replication to complete. While the simulations might in rare cases generate unrelasitic situations, for example if a large initiation adder is combined with a small division adder leading to premature division, those clashes seem rare enough to not affect the global statistical behavior of the model. Naturally the model would break down and these controls would need to be explicitly included in the case where cells are subject to stress conditions where these control mechanisms act as fail-safes e.g. to ensure that division is delayed if DNA repair is needed. In the course of our study, new research ***Si et al. (2019***) proposed that division and replication cycles are only seemingly connected, and used perturbation methods to drive cells to states where the uncoupling is revealed. While such perturbation studies are very informative, more work is needed to understand to what extent hidden compensatory mechanisms might be at play e.g. when affecting DnaA or FtsZ expression. Also, that study focuses exclusively on a model which explicitly enforces various correlations between variables unlike our model which naturally produces such relations. It would thus be a worthwhile future endeavour to test our simpler model on such perturbed growth conditions, a task beyond the purpose of this study, which tries to clarify the normal growth case.

Interestingly, a double-adder mechanism similar to the one that we propose here has been shown to explain cell-cycle control in mycobacteria (***Logsdon et al., 2017***). These mycobacteria have a much more complex behavior than *E. coli*, in particular characterized by a strong asymmetry between daughter cells and a growth rate almost an order of magnitude smaller than that of *E. coli*. Despite those important differences, it was shown that mycobacterial cell cycles exhibit adder behavior for both division and replication starting at initiation, in a manner highly similar to our observations in *E. coli*. This suggests that the mechanism connecting replication and division must be quite fundamental and independent of the specifics of available genes and their expression.

Although the single-cell observations provide clear indications of which variables are most likely to be directly involved in the cell cycle control, they of course do not indicate the underlying molecular mechanisms. However, it is not hard to speculate about possible molecular mechanisms that could implement the double-adder behavior. As others have pointed out previously (***Ho and Amir, 2015***), an adder for the regulation of replication initiation can be easily implemented at the molecular level by having a “sensor” protein that builds up at each origin, and that triggers replication initiation whenever a critical mass is reached at a given origin. If this sensor protein is additionally homeostatically controlled such that its production relative to the overall protein production is kept constant, then the average volume per origin will also be kept constant across conditions.

It is more challenging to define a molecular system that can implement the second adder that controls division. The main challenge is that this adder does not run throughout the entire cell cycle, but only between replication initiation and division. It is well known that division is driven by the polymerization of the FtsZ ring, which includes a host of other FtsZ-ring associated proteins, and its progressive constriction. It might seem simplest to assume that the division adder could be implemented directly through FtsZ production, again in the logic of the regulated “sensor” mentioned above. However, this would require FtsZ to be produced and accumulating at the division sites only from replication initiation to cell division. Although this is conceivable, i.e. it is known that FtsZ and other division proteins are heavily regulated at several levels (***Dewar and Dorazi, 2000***) and that especially in slow growth conditions its concentration varies during the cell cycle (***Männik et al., 2018***), it is hard to imagine how this model could work under fast growth conditions in which there are overlapping rounds of replication such that FtsZ would be constantly expressed. Moreover recent data (***Si et al., 2019***) rather suggest that FtsZ concentration is constant during the cell cycle.

Alternatively, rather than FtsZ production, Ftsz polymerization could be regulated. One remarkable observation that is well known within the field (***Lau et al., 2004***; ***Nielsen et al., 2006***) and that we also observe in our data (see ***Figure Supplement 1***), is that origins always occupy the position of future division sites (mid-cell, 1/4 and 3/4 positions etc.) when replication is initiated. This observation not only suggests that, at replication initiation, some local molecular event occurs that will eventually trigger division at the same site, but it is also remarkably consistent with the idea of an adder running only between replication initiation and division. One long-standing idea that is consistent with these observations is that some molecular event that occurs during replication initiation triggers the start of FtsZ ring formation, and that the timing from initiation to division is controlled by the polymerization dynamics of the FtsZ ring (***Weart and Levin, 2003***). At the molecular level, the common triggering of initiation and polymerization might be explained by the accumulation of acidic phosholipids in the cell membrane precisely at future division sites (***Renner and Weibel, 2011***) where they probably interact with components of the division machinery. At the same time those lipids are known to play a role in promoting replication by rejuvenating the initiator protein DnaA-ADP into DnaA-ATP (***Saxena et al., 2013***), and might therefore be a “hub” coordinating the two cycles. Finally, it remains to be explained how FtsZ polymerization or pole building could result in an adder beahviour. For that purpose, future experiments should focus on combining the type of information collected in this study and detailed measures of the dynamics of FtsZ-ring assembly and constriction as done in ***Coltharp et al. (2016***).

## Methods and Materials

### Bacterial strains and media

All strains are derived from the K-12 strain BW27378, a Δ(araH-araF)570(::FRT) derivative of the Keio collection background strain (***Baba et al., 2006***) obtained from the Yale Coli Genetic Stock Center. This strain was further modified by *λ*-Red recombination (***Datsenko and Wanner, 2000***) and P1 transduction to result in ΔaraFGH(::FRT), ΔaraE(::FRT), ΔlacIZYA(::FRT). A 250 lacO repeats FROS array with chloramphenicol resistance was inserted close to the origin of replication in the *asnA* gene by *λ*-Red recombination and P1 transduction. The lacO-CmR array was derived from the original plasmid pLau43 (***Lau et al., 2004***) by replacing the kanamycin resistance and a series of operators on both sides of it with the CmR gene. For visualization of the array, LacI-mVenus was expressed from the plasmid pGW266, derived from the original FROS plasmid pLAU53 (***Lau et al., 2004***) from which the tetR construct was removed and the lacI-CFP replaced by lacI-mVenus. For the experiment analyzed automatically the same stain carried in addition the plasmid pGW339 expressing FtsZ-mVenus under the control of the araBAD promoter using 0.002% arabinose for induction. Expression is tighly controlled by using the approache proposed in ***Morgan-Kiss et al. (2002***).

All experiments were done using M9 minimal media supplemented with 2mM MgSO4, 0.1mM CaCl2, and sugars (0.2 % for glucose and 0.2 % for glycerol). In one experiment, the media was supplemented with 8 amino acids at a concentration of 5 μg/ml: Threonine, Aspagrinine, Methionine, Proline, Leucine, Tryptophane, Serine, Alanine. All experiments were carried out at 37 °C.

### Microfluidic device fabrication

Mother Machine experiments were performed using the Dual Input Mother Machine (DIMM) microfluidic design which has been described elsewhere (***Kaiser et al., 2018***) and is freely available online (https://metafluidics.org/devices/dual-input-mother-machine/); since no change of conditions was intended during experiments, the same media was flown at both inputs.

Several microfluidics masters were produced using soft lithography techniques by micro-resist Gmbh; two masters with regular growth channels of suitable size (0.8 μm width × 0.9 μm height for growth in glycerol, and 1 μm width × 1.2 μm height for growth in glucose) were used for all experiments.

For each experiment, a new chip was produced by pouring PDMS (Sylgard 184 with 1:9w/w ratio of curing agent) on the master and baking it for 4 h or more at 80°C. After cutting the chip and punching inlets, the chip was bonded to a #1.5 glass coverslip as follows: the coverlsip was manually washed in water and soap, rinsed in isopropanol then water; the chip cleaned from dust using MagicTape, rinsed in isopropanol then water; surfaces were activated with air plasma (40 sec at 1500 μm of Hg) before being put in contact; the assembled chip was cooked 1 h or more at 80°C.

Before running the experiment, the chip was primed and incubated1h at 37°C using passivation buffer (2.5 mg/mL salmon sperm DNA, 7.5 mg/mL bovine serum albumin) for the mother machine part and water for the overflow channels.

## Experiment setup and conditions

Bacteria were stored as frozen glycerol stocks at −80 °C and streaked onto LB agar plates to obtain clonal colonies. Overnight precultures were grown from single colonies in the same growth media as the experiment. The next day, cells were diluted 100-fold into fresh medium and harvested after 4-6 h.

The experimental apparatus was initialized, pre-warmed and equilibrated. Media flow was controlled using a pressure controller and monitored with flow-meters, set to run a total flow of ≈1.5 μL/ min (corresponding to a pressure of ≈1600 mbar).

The primed microfluidic chip was mounted, connected to media supply and flushed with running media for 30 min or more to rinse passivation buffer. The grown cell culture was centrifuged at 4000×g for 5 min, and the pellet re-suspended in a few μL supernatant and injected into the device from the outlet using the pressure controller. To facilitate the filling of growth channels by swimming and diffusing cells, the pressure was adjusted in order to maintain minimal flow in the main channel (loading time 40min).

After loading, bacteria were incubated during 2 h before starting image acquisition. Every 3 min, phase contrast and fluorescence images were acquired for several well-separated positions in parallel.

### Microscopy and image analysis

An inverted Nikon Ti-E microscope, equipped with a motorized xy-stage and enclosed in a temperature incubator (TheCube, Life Imaging Systems), was used to perform all experiments. The sample was fixed on the stage using metal clamps and focus was maintained using hardware autofocus (Perfect Focus System, Nikon). Images were recorded using a CFI Plan Apochromat Lambda DM ×100 objective (NA 1.45, WD 0.13 mm) and a CMOS camera (Hamamatsu Orca-Flash 4.0). The setup was controlled using μManager (***Edelstein et al., 2014***) and timelapse movies were recorded with its Multi-Dimensional Acquisition engine. Phase contrast images were acquired using 200 ms exposure (CoolLED pE-100, full power). Images of mCherry fluorescence were acquired using 200ms exposure (Lumencor SpectraX, Green LED at 33% with ND4) using a Semrock triple-band emission filter (FF01-475/543/702-25).

Image analysis was performed using MoMA (***Kaiser et al., 2018***) as described in its documentation (https://github.com/fjug/MoMA/wiki). Raw image datasets were transferred to a centralised storage and preprocessed in batch. Growth channels were chosen randomly after discarding those where cell cycle arrest occurred in the mother cell, and curated manually in MoMA. An exponential elongation model was then fitted to each cell cycle, and cycles presenting large deviations were discarded (1-3% of each experiment).

For the automated origin detection and tracking we used custom Python code. Spots were detected following the method proposed in ***Aguet et al. (2013***). Briefly, amplitude and background are estimated for each pixel using a fast filtering method and a spot model corresponding to the optical setup. Among the local maxima found in the amplitude estimates, spots are then selected using a statistical test based on the assumption that background noise is Gaussian. To track spots we used the trackpy package ***Allan et al. (2018***). We kept only cell cycles where origin tracks behaved in a biologically reasonable way, *i.e.* one track splitting in two, splitting in four *etc*. The time of initiation was assigned as the first time point where a track splits into two. For the manual analysis of the other experiments, the frame showing origin splitting was selected manually.

Using the the timing of origin splitting, the corresponding cell length could be determined. All further variables like *dL* or *d*Λ_*ib*_ are deduced from the primary variables. For the decomposition analysis, a pseudo-cell cycle was created by concatenating the mother cell cycle from initiation to division with the daughter cell cycle from birth to initiation. The growth rate *α* for this pseudo-cell cycle was again obtained by fitting an exponential growth model. All the growth lanes corresponding to a given conditions were then pooled to generate the various statistics shown in this article.

The entire analysis pipeline is available as Python modules and Jupyter Notebooks on Github (https://github.com/guiwitz/DoubleAdderArticle).

### Simulations

The numerical implementation of the model described in ***Figure 4*** and used in ***Figure 5*** requires several parameters for each individual cell cycle. To generate those, the following distributions were extracted from experimental data, and if needed their means and variances were obtained by a fitting procedure:

- The growth rate distributions *P* (*λ*).
- The growth rate correlation from mother to daughters.
- The length distributions of the two adder processes *P* (*d*Λ_*ib*_) and *P* (*d*Λ_*if*_).
- The distributions of length ratios between sister cells to account for imprecision in division placement *P* (*r*).

For the simulation, a series of 500 cells is initialized with all required parameters: initial length *L*_0_ taken from the birth length distribution, *λ* = *P* (*λ*), number of origins *n*_*ori*_ = 1, and the two adders *d*Λ_*ib*_ = *P* (*λ*) and *d*Λ_*if*_ = *P* (*d*Λ_*if*_)) whose counters are starting at 0. The exact initialization is not crucial as the system relaxes to its equilibrium state after a few generations. Cells are then growing incrementally following an exponential law, and the added length is monitored. Every time the cell reaches its target *d*Λ_*if*_, the number of origins doubles and a new initiation adder is drawn from *P* (*d*Λ_*if*_). Every time the cell reaches its target *d*Λ_*ib*_ the cell 1) divides into two cells using a division ratio drawn *P* (*r*), 2) the number of origins per cell is divided by two, 3) a new division adder is drawn from *P* (*d*Λ_*ib*_), and finally 4) a new growth rate is drawn from *P* (*λ*). Each simulation runs for 30h in steps of 1min. In the end, the cell tracks resulting from the simulation are formatted in the same format as the experimental data, and follow the same analysis pipeline. The code is available on Github.

## Acknowledgments

### Additional information

This study was funded through the SNSF Ambizione grant PZ00P3-161467 to GW and the SNSF grant 31003A-159673 to EvN.

## Appendix 1 Experiments statistics

**Table 1.** Statistics for all expderiments. Glycerol auto is that dataset analyzed automatically, while Glycerol is the one analyzed manually. *r* stands for Pearson correlation, and the *m* − *d* superscript indicates a mother-daughter correlation. The doubling time (1/*λ*) is obtained by fitting the distribution of growth rates with a log-normal distribution.

| Experiment | Discarded % | # cell cycles | $1/\lambda[\text{min}]$ | Adder $r$ | $\lambda^{m-d} r$ | $L_b^{m-d} r$ |
| --- | --- | --- | --- | --- | --- | --- |
| Glycerol auto | 3.3 | 3070 | 86.0 | -0.10 | 0.33 | 0.45 |
| Glycerol | 2.1 | 810 | 89.0 | -0.07 | 0.42 | 0.58 |
| Glucose | 2.1 | 1035 | 53.0 | -0.04 | 0.47 | 0.66 |
| Glucose +AA | 2.4 | 1159 | 41.0 | -0.12 | 0.36 | 0.48 |

## Appendix 2 Other models

In this article we have shown that models relying on the concept of initiation mass, as well as those involving a constant timer from initiation to division are incompatible with measurements. Still, those models are able to reproduce a wide range of experimental measurements, and we wanted to understand where they would break. We give here two examples of such an analysis. In the first case we tried to reproduce the model proposed in ***Wallden et al. (2016***). This model assumes that cells initiate replication around a specific initiation mass length *L*_*i*_ and then grow for an amount of time depending on growth rate *T*_*CD*_(*μ*) before dividing (1A). In panels B and C of 1 we show that we are successfully reproducing the model used e.g. in Figure 6 of ***Wallden et al. (2016***). The histogram of the number of origins at birth shown in 1D shows a clear failure of the model where cells in slow growth conditions are all born with an ongoing round of replication in contradiction with experimental data (see e.g. Figure 3 of ***Wallden et al. (2016***)).

**Figure 1.**
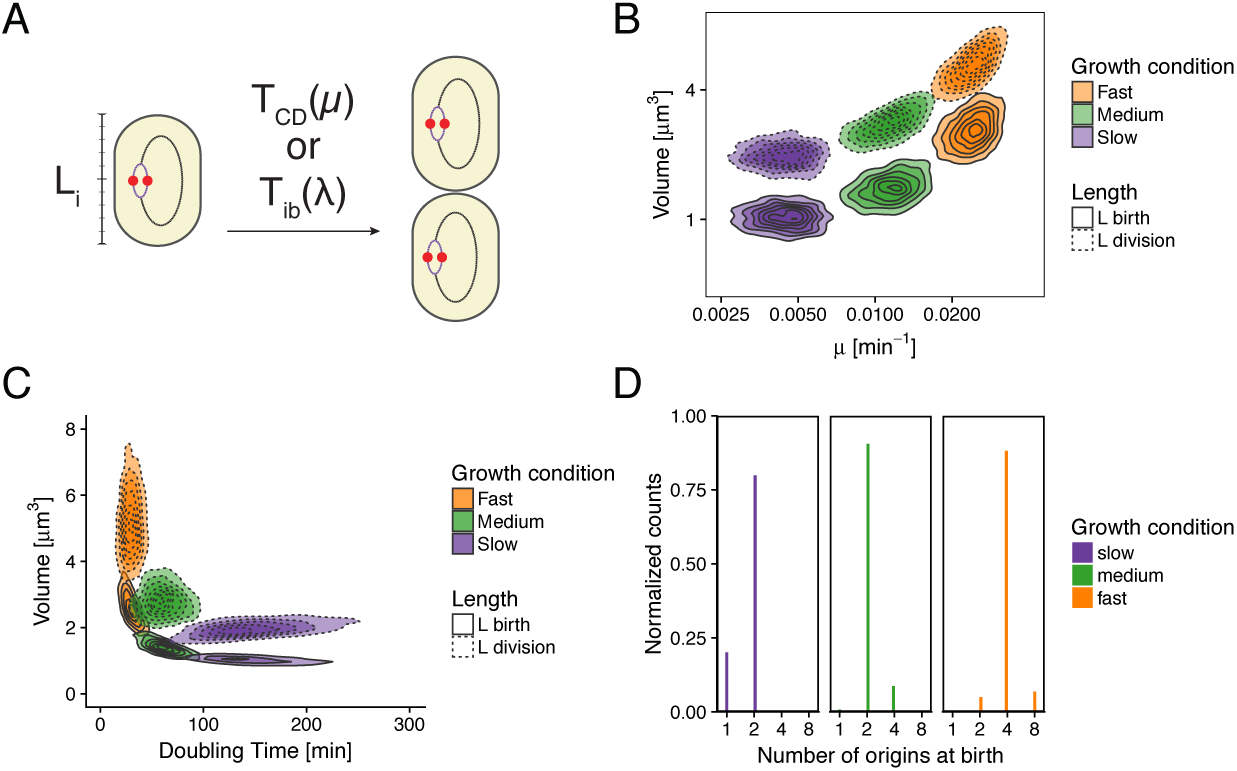
Re-implementation of the model proposed in ***Wallden et al. (2016***) for three growth conditions. A. Cells initiation at length *L*_*i*_ and grow fora time *T*_*CD*_(*μ*) before dividing. B. Cell volume at birth and division as a function of growth rate. C. Cell volume at birth and division as a function of generation time. D. Distributions of the number of origins at birth.

The second model we are investigating here has been recently proposed by ***Micali et al. (2018b)***. It uses an inter-initiation adder for replication regulation, and combines it with a classical adder (birth to division) without coupling those two regulation systems together explicitly. We simulated such a model with the added constraint that division can only occur if at least two origins are present in the cell. The results are shown in Fig.2. The model surprisingly reproduces most of the features of the experimental data with one exception: the initiation to division variable *d*Λ_*ib*_ is clearly not anymore an adder. This can be trivially explained: as the two mechanisms are uncoupled, an initiation at a large size automatically leads to a small *d*Λ_*ib*_ on average while an early initiation at small size leads to a large *d*Λ_*ib*_ on average.

**Figure 2.**
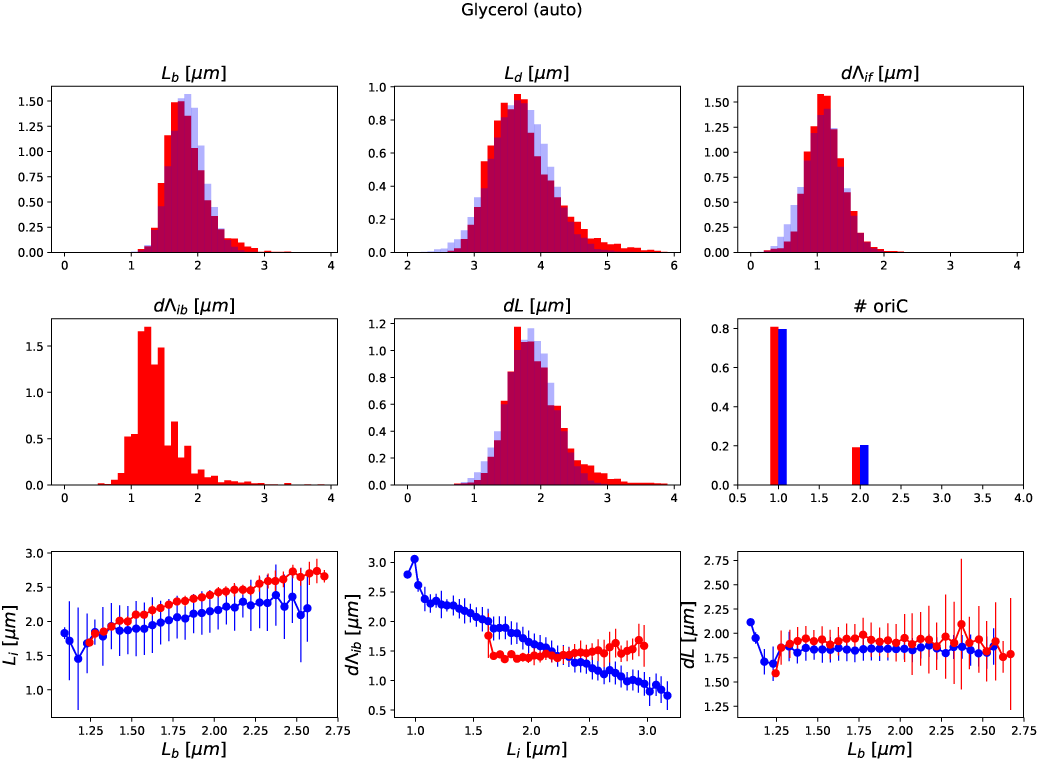
Comparison of experimental distributions and correlations with a model combining an inter-initiation adder and a classic adder *dL* = *L*_*d*_ − *L*_*b*_

## Appendix 3 Decomposition: the classic adder model

In order to illustrate the functioning of the our decomposition approach, we apply it to the familiar case of the classic division adder model. By considering all possible combinations of standard cell cycle variables and estimating their independence, we find that the decomposition offering the most independent set of variables corresponds to the classic adder model defined by *L*_*b*_, *dL* and *λ*, as can be seen in 1.

**Figure 1.**
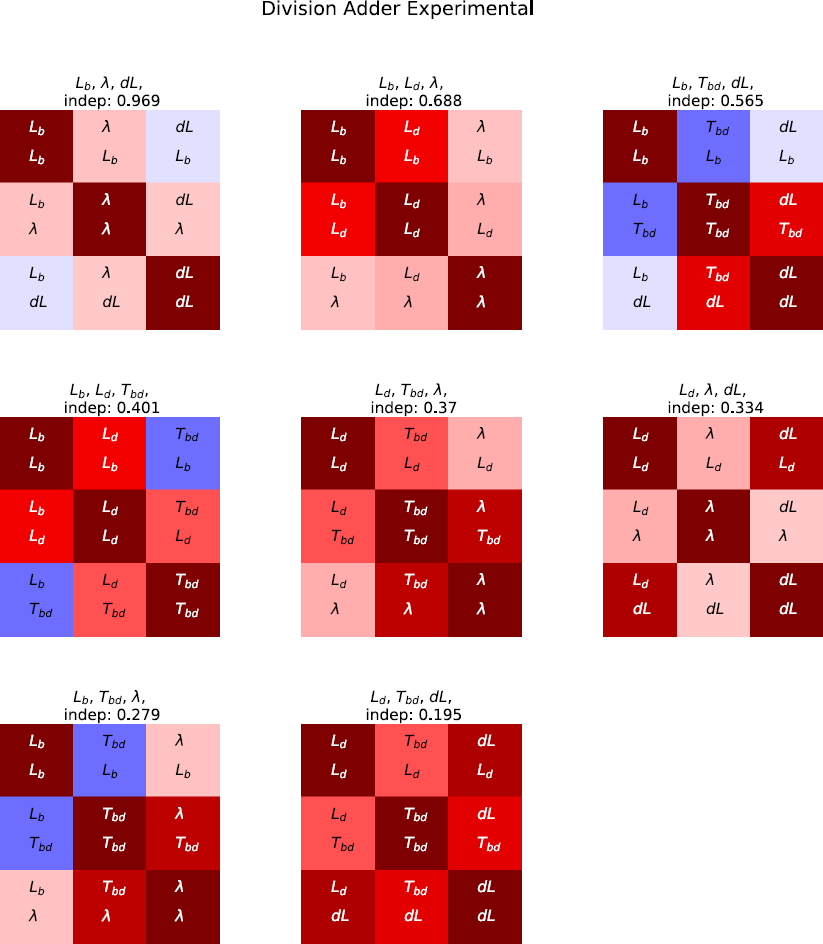
All possible decompositions for the division cycle variables ranked from best (top left) to worst (bottom right) in terms of variable independence.

---

**Figure 1– Figure supplement 1.**
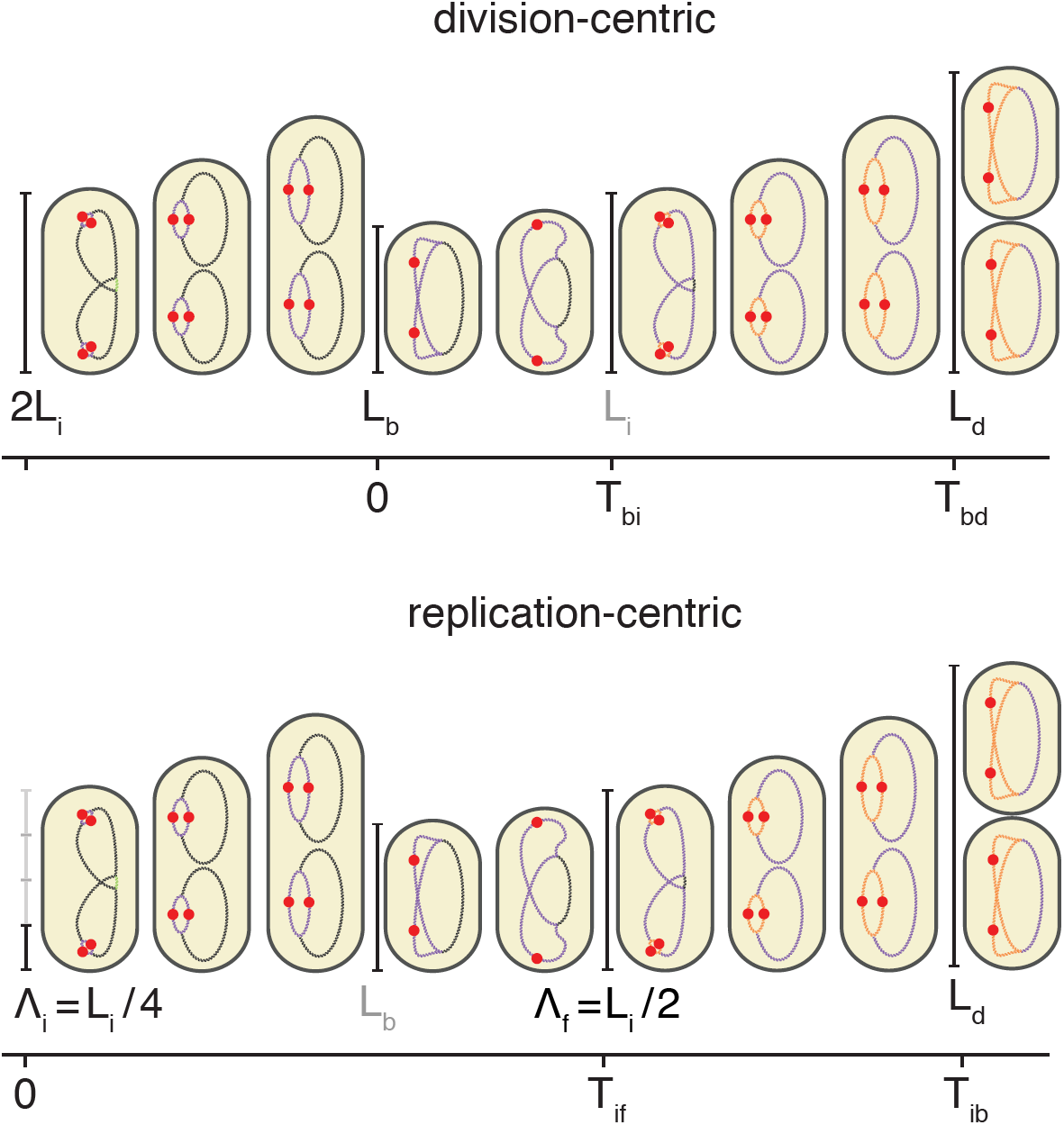
Schema of the cell cycle and variable definitions for the case of fast growth with overlapping replication cycles

**Figure 4– Figure supplement 1.**
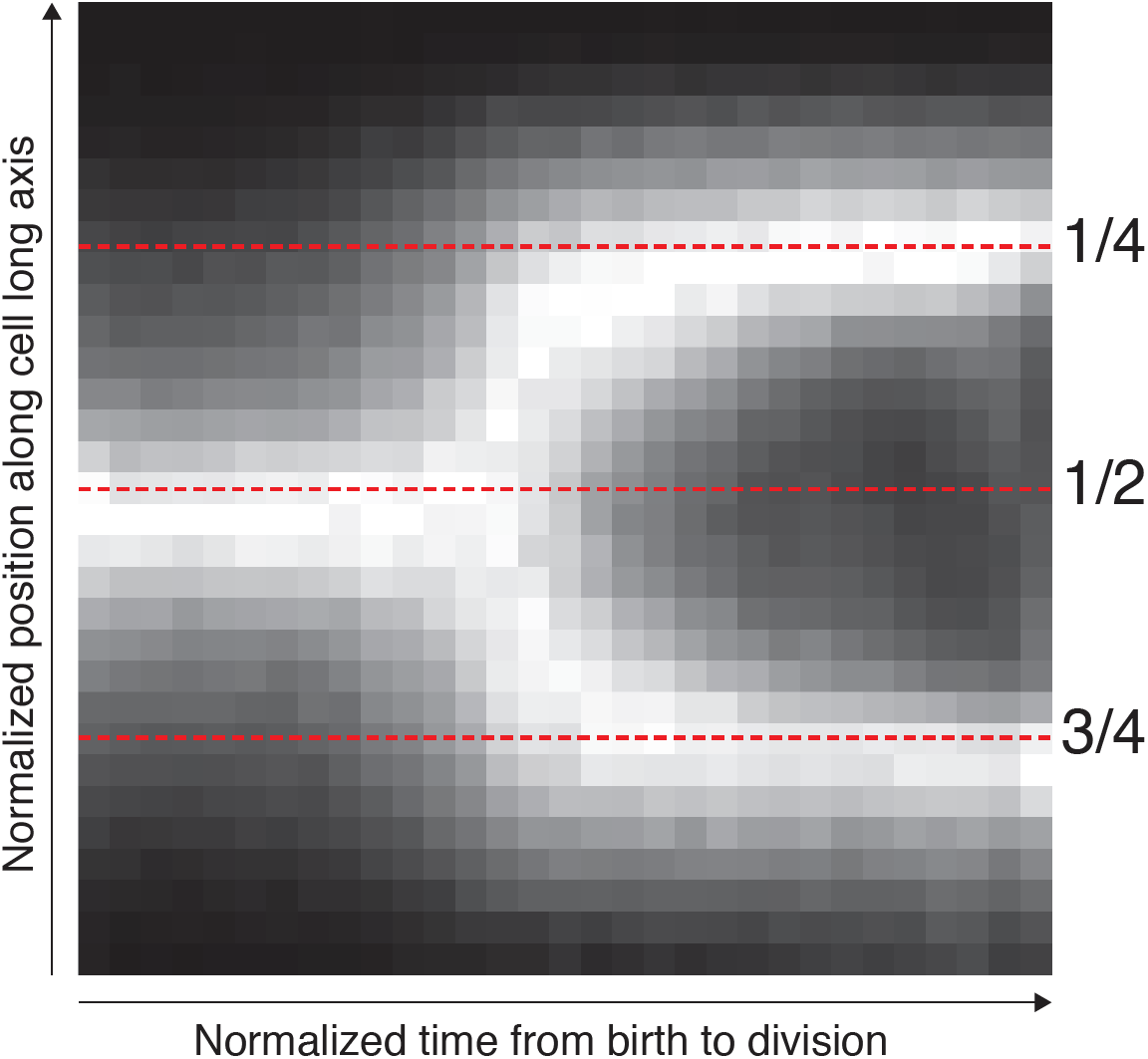
The position along the cell axis and the cell-cycle time of all detected spots were collected. The longitudinal position was scaled with cell length to indicate the relative position in the cell. The cell cycle time was normalized between 0 and 1. The 1 shows as 2D histogram of these space-time data. The intensity of each time point (columns) has been normalized. The mid-cell and quarter-cell (mid-cell of daughter cell) positions are indicated with dotted lines.

**Figure 5– Figure supplement 1.**
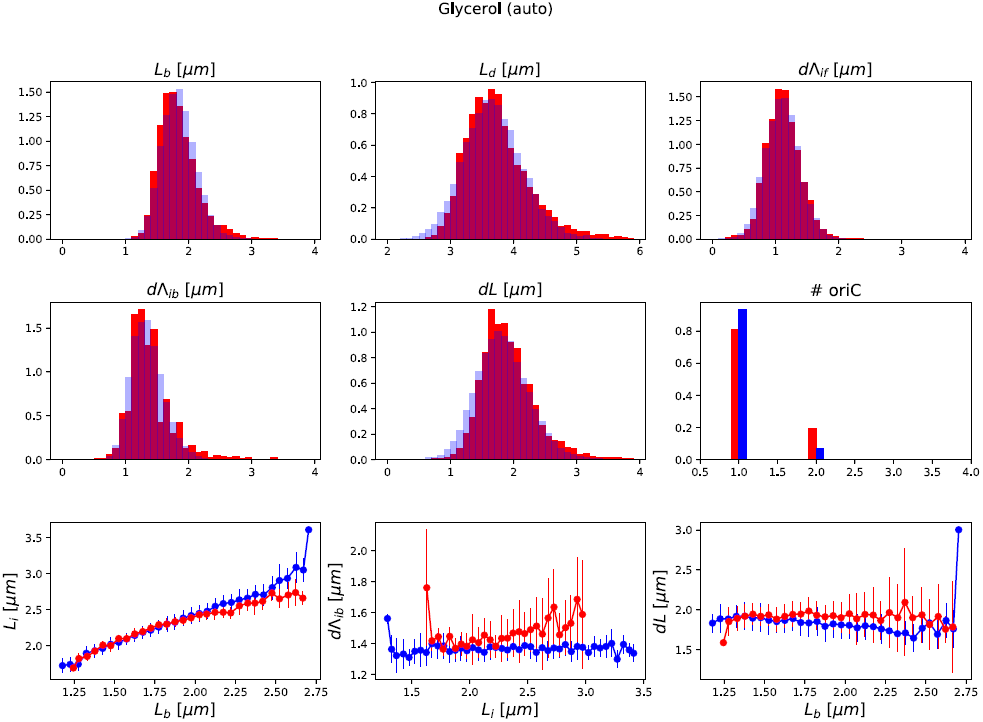
Detailed comparisons between experiments and simulations for M9+glycerol condition (with automated origin tracking).

**Figure 5– Figure supplement 2.**
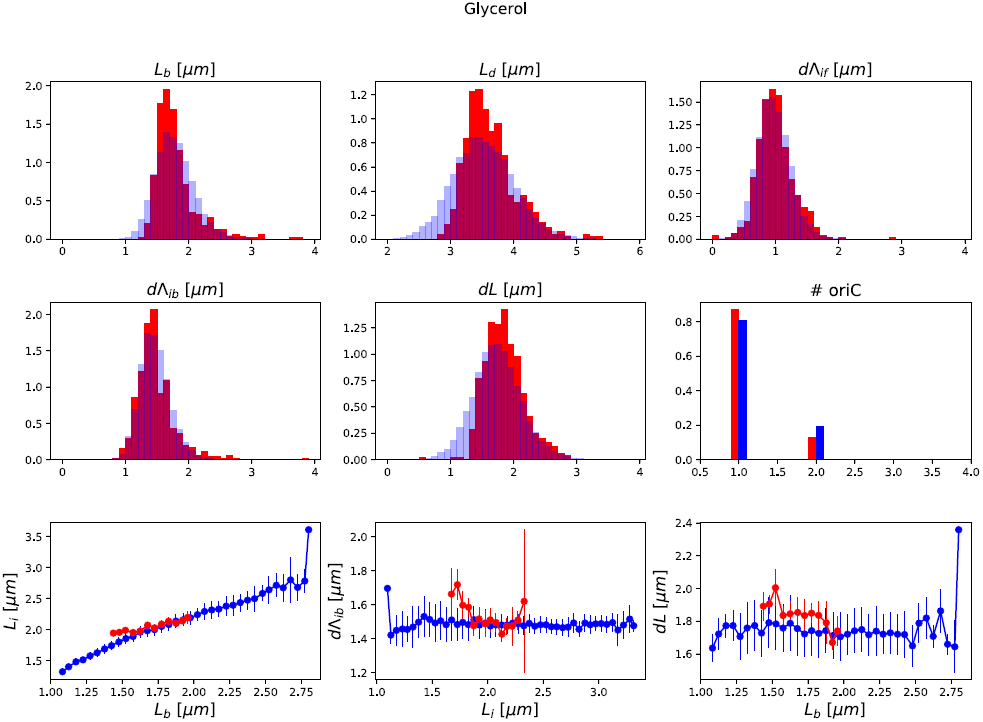
Detailed comparisons between experiments and simulations for M9+glycerol condition (with manual origin tracking).

**Figure 5– Figure supplement 3.**
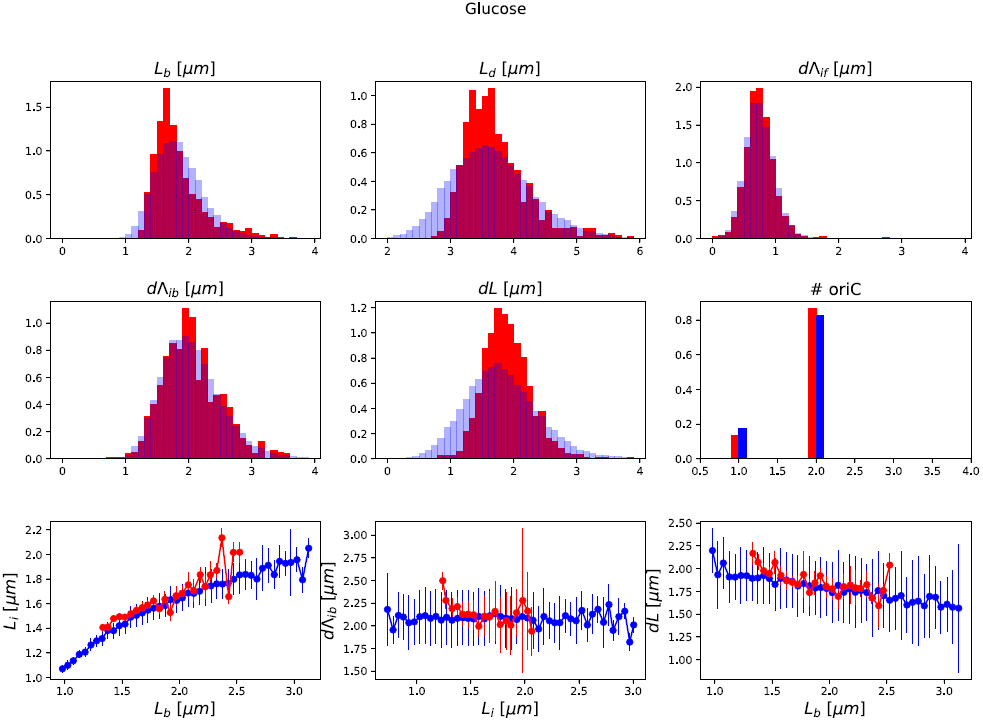
Detailed comparisons between experiments and simulations for M9+glucose condition (with manual origin tracking).

**Figure 5– Figure supplement 4.**
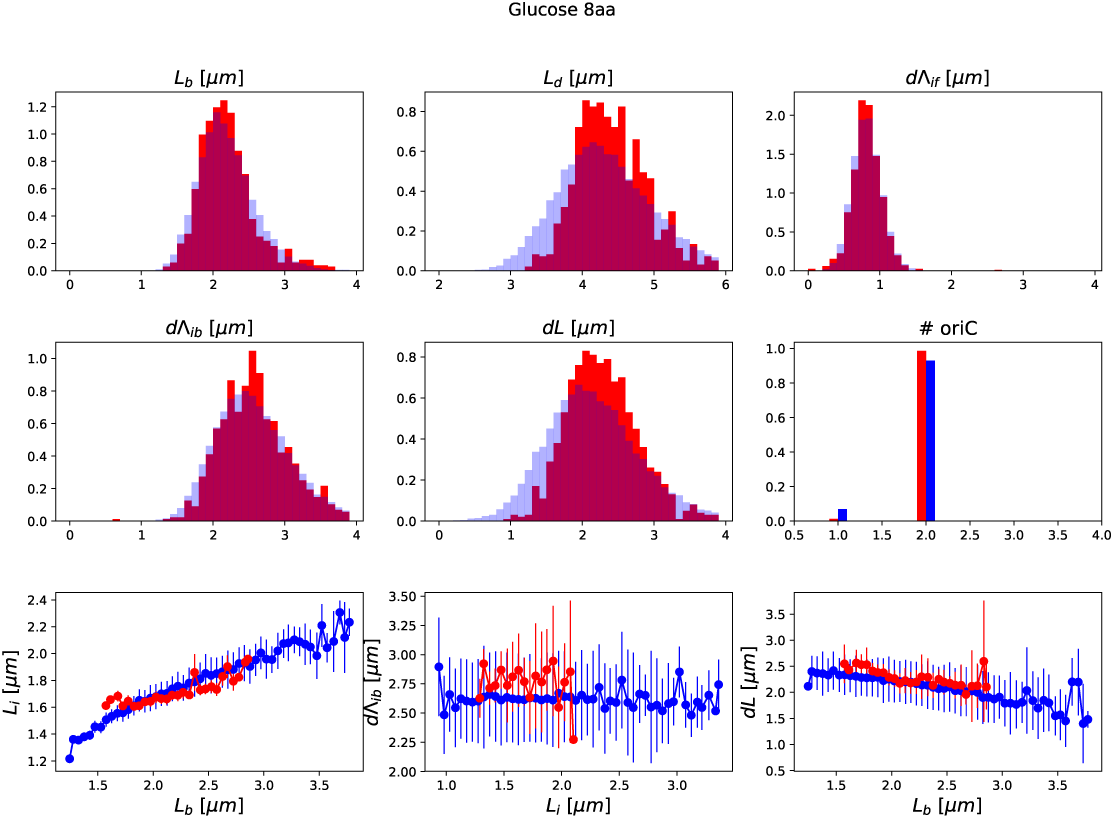
Detailed comparisons between experiments and simulations for M9+glucose+8a.a. condition (with manual origin tracking).

**Figure 5– Figure supplement 5.**
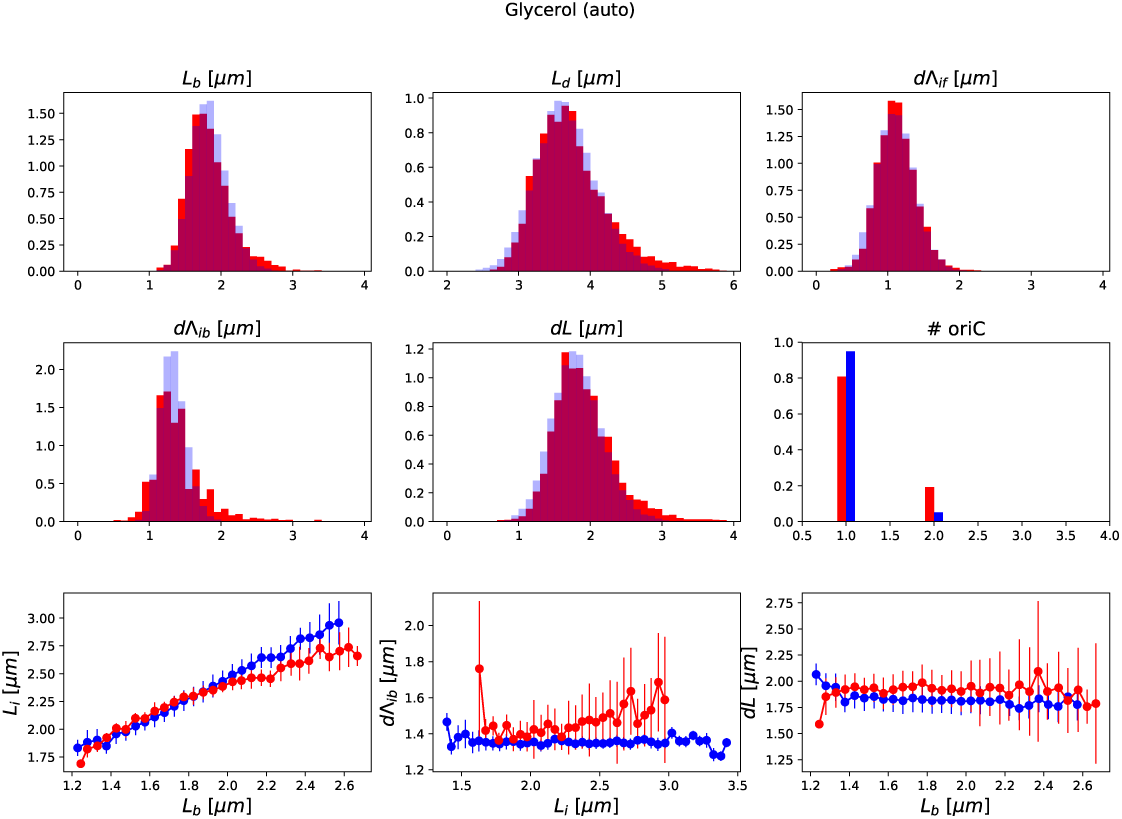
Focusing on the large dataset (with automated origin tracking) for which we have the most accurate measures, we note that *dL* shows a slight deviation from adder behavior. As shown here, we found that this could be corrected by slightly reducing the variance level of the division adder distribution (to 70% of its original value). As the initiation measurement is made imprecise for experimental (*e.g.* acquisition rate) and biological (*e.g.* variable cohesion of origins), it is reasonable to assume that we overestimate the variance of that parameter.

**Figure 6– Figure supplement 1.**
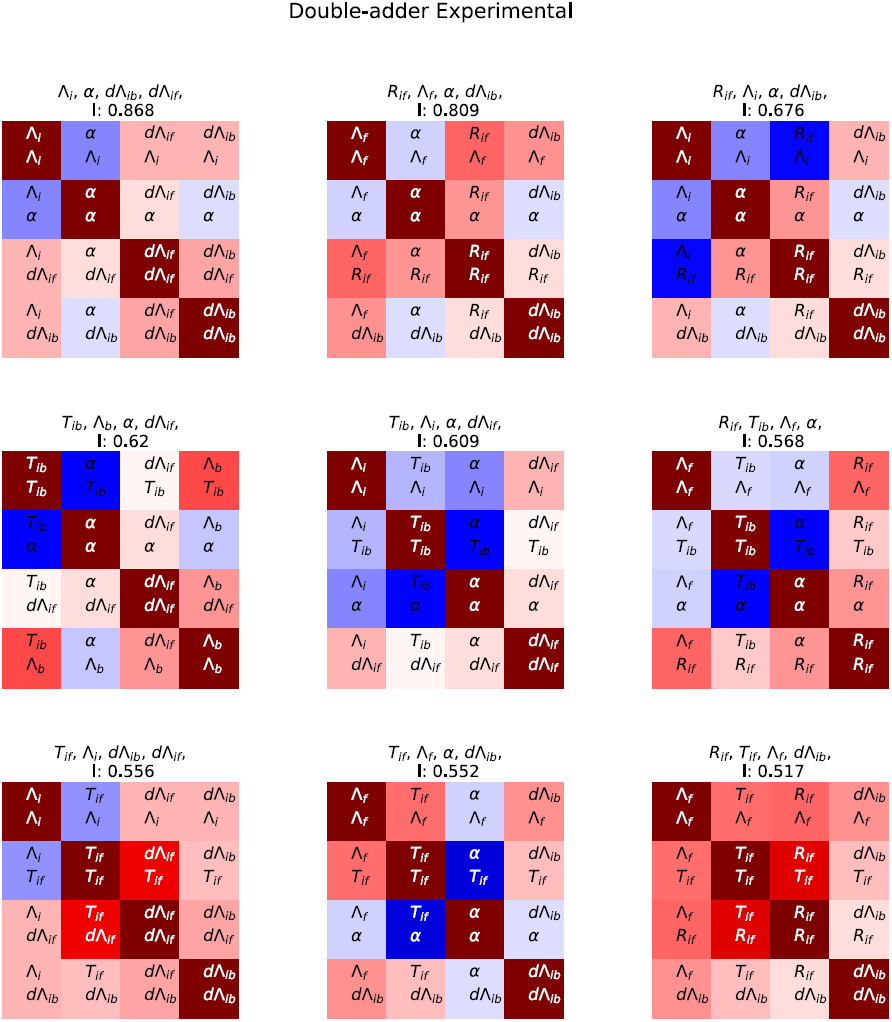
In complement to the best decomposition shown in Fig.6, we show here details for the first nine best decompositions for the experimental data.

**Figure 6– Figure supplement 2.**
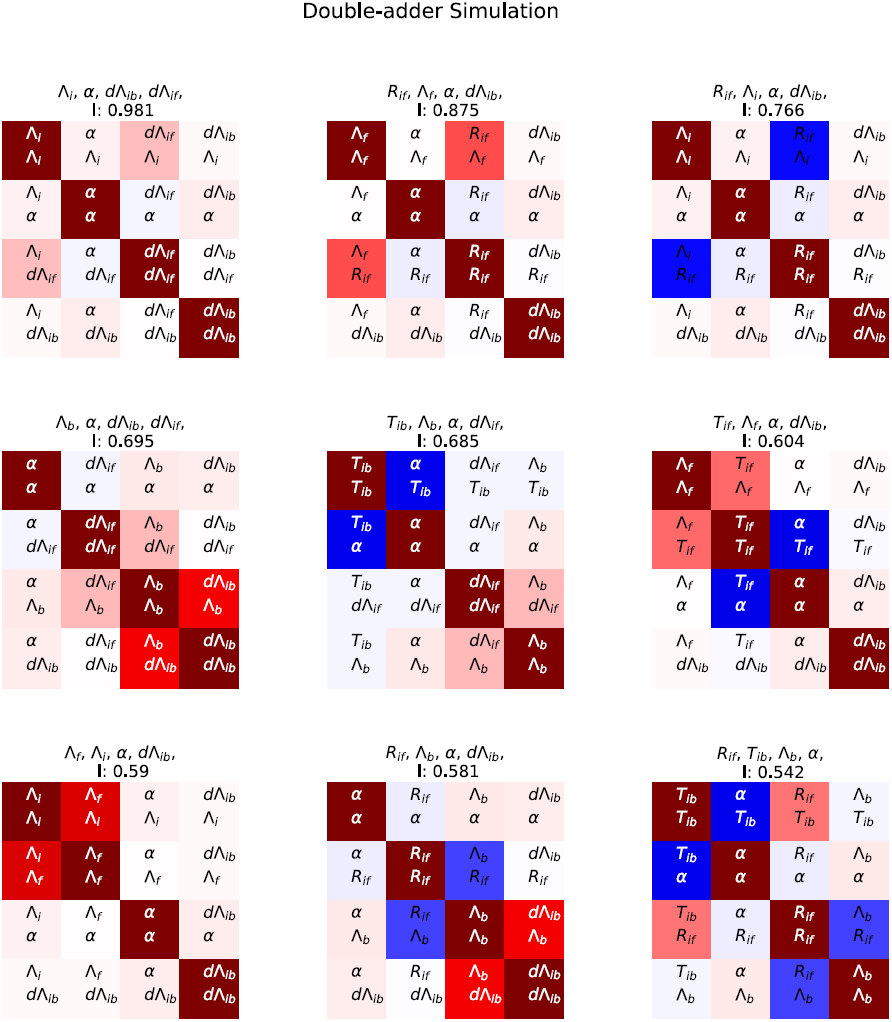
In complement to the best decomposition shown in Fig.6, we show here details for the first nine best decompositions for the simulation data.

**Figure 6– Figure supplement 3.**
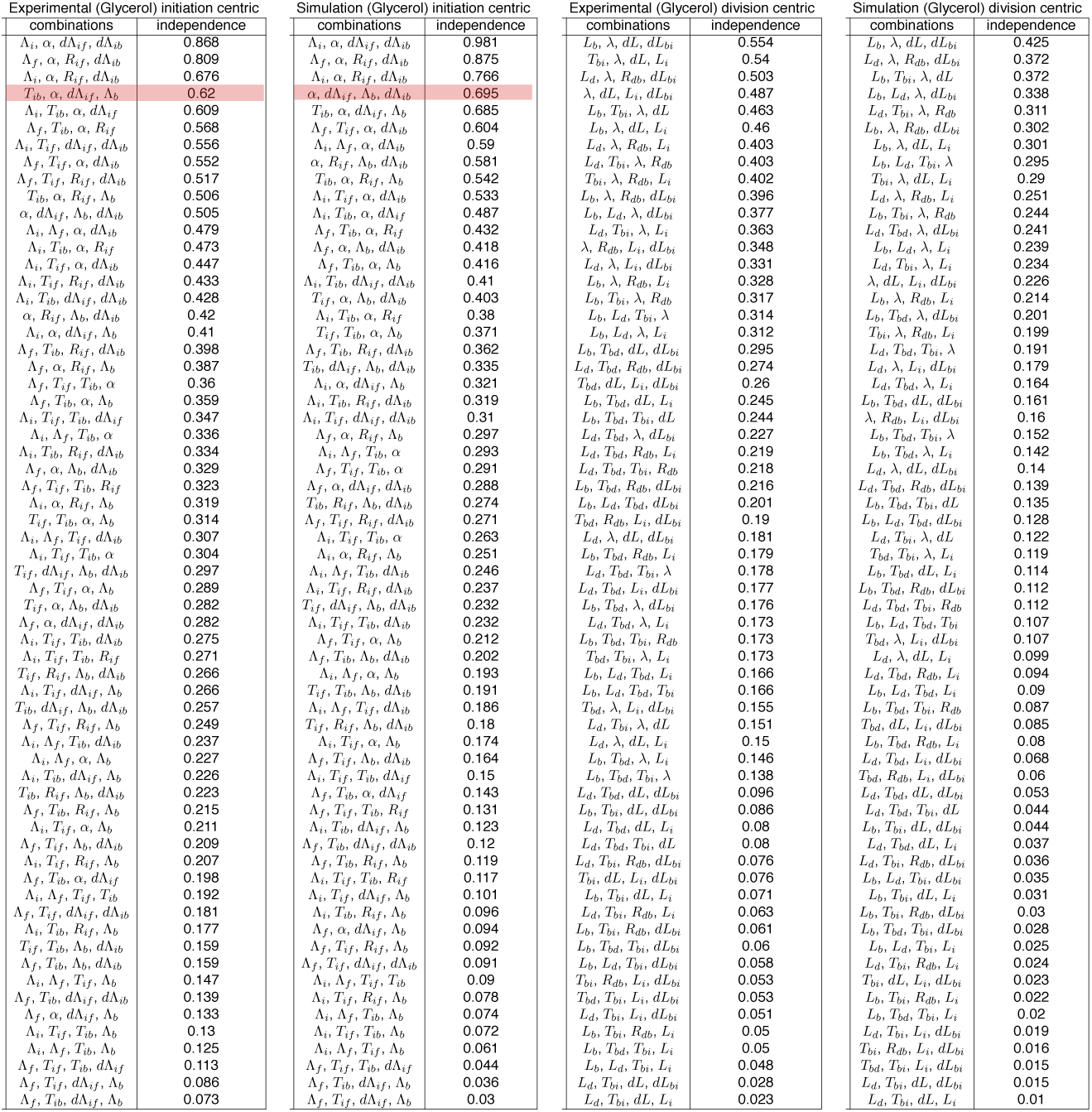
All possible decompositions for both experimental and simulation data are shown ranked from most to least independent.

